# Epidemiological and ecological modelling reveal diversity in upper respiratory tract microbial population structures from a cross-sectional community swabbing study

**DOI:** 10.1101/099069

**Authors:** Abigail L. Coughtrie, Denise E. Morris, Rebecca Anderson, Nelupha Begum, David W. Cleary, Saul N. Faust, Johanna M. Jefferies, Alex R. Kraaijeveld, Michael V. Moore, Mark A. Mullee, Paul J. Roderick, Andrew Tuck, Robert N. Whittaker, Ho Ming Yuen, C. Patrick Doncaster, Stuart C. Clarke

## Abstract

Respiratory tract infections (RTI) are responsible for over 4 million deaths per year worldwide with pathobiont carriage a required precursor to infection. Through a cross-sectional community-based nasal self-swabbing study we sought to determine carriage epidemiology for respiratory pathogens amongst bacteria (*Streptococcus pneumoniae, Haemophilus influenza, Moraxella catarrhalis, Staphylococcus aureus, Pseudomonas aeruginosa* and *Neisseria meningitidis*) and viruses (RSV, Influenza viruses A and B, Rhinovirus/Enterovirus, Coronavirus, Parainfluenza viruses 1-3 and Adenovirus (ADV)). Carriage of bacterial and viral species was shown to vary with participant age, recent RTI and the presence of other species. The spatial structure of microbial respiratory communities was less nested (more disordered) in the young (0-4 years) and those with recent RTI. Species frequency distributions were flatter than random expectation in young individuals (X^2^ = 20.42, *p* = 0.002), indicating spatial clumping of species consistent with facilitative relationships amongst them. Deviations from a neutral model of ecological niches were observed for samples collected in the summer and from older individuals (those aged 5-17, 18-64 and ≥65 years) but not in samples collected from winter, younger individuals (those aged 0-4 years), individuals with recent RTI and individuals without recent RTI, demonstrating the importance of both neutral and niche processes in respiratory community assembly. The application of epidemiological methods and ecological theory to sets of respiratory tract samples has yielded novel insights into the factors that drive microbial community composition, such as seasonality and age, as well as species patterns and interactions within the nose.

## INTRODUCTION

Respiratory tract infections (RTI) remain a significant cause of mortality worldwide, with 2.8 million deaths, from an estimated four million per annum, resulting from lower RTI and a further 3,000 deaths resulting from upper RTI in 2010 (1). An estimated six million individuals in the UK visit their general practitioner each year regarding a RTI, posing a substantial burden on the health service (2).

Microbial carriage of pathogens in the upper respiratory tract is a precursor to RTI, meningitis and sepsis and can also facilitate the transmission of microbes between individuals. Carriage is a dynamic process involving fluctuations of species and strains (3). The interactions and interplay between bacterial and viral species within the respiratory tract is thought to affect the shift from asymptomatic carriage to serious invasive illness (4). Such interactions will be driven by species presence and diversity of the resident communities of the respiratory tract. Here diversity in species richness and abundance is thought to offer protection from invading pathogens and overgrowth of a single species (5, 6). It is likely that the dynamics of community structure may impact on infection risk. Species diversity can be measured in the presence of different species within the respiratory tract (inter-species diversity) or the presence of different bacterial strains and types of a single species (intraspecies). The former can be assessed using conventional culture methods or molecular techniques such as real-time polymerase chain reaction (PCR) and 16S rRNA gene sequencing (7, 8). Intra-species analysis gives insight into the distribution of different types of a species that may display differential capacity for disease. It requires techniques such as serotyping and multi-locus sequence typing (MLST) (9–12), or expansions of the MLST methodology such as whole-genome MLST (wgMLST) (13) and ribosomal MLST (rMLST) (14).

The processes that sustain biodiversity may involve species occupying more or less distinct ecological niches, or they may involve neutral processes of stochastic zero-sum dynamics amongst more or less identical species (15). Niche dynamics describe the distinct nutritional and environmental influences that determine the occurrences and abundances of different species (16, 17). Neutral dynamics describe stochastic influences on species abundance and diversity, such as birth, death, immigration, dispersal and speciation (5, 18). These alternative dynamics have been much studied in terms of species distribution and community assembly patterns for plant-insect (19), plant-fungi (20) and parasite-fish (21) interactions. They have not, to our knowledge, been applied to respiratory tract microbial ecology.

Here, we aim to understand how bacterial and viral species maintain diversity within the respiratory tract, the nature of their interactions and the factors influencing community assembly and colonisation. To our best knowledge this is the first study that aims to discern multi-species epidemiology using a broad cross-sectional community cohort with the added dimension of seasonality. This was achieved using culture-based techniques, molecular methods, mathematical modelling and the application of ecological models.

## METHODS

### Sample Collection and Analysis

Nose swab samples were collected as part of a large population-based cross-sectional carriage study undertaken with two separate time-points, May to August 2012 (late spring/summer) and February to April 2013 (winter/early spring), following protocols described in reference (22). Briefly this involved the collection of nose swabs from the two time-points were collected from different individuals from twenty general practitioners (GP) practices from the South East hub of the Wessex Primary Care Research Network. Figure 1 depicts the structure of the study. Swabs were immersed into skimmed milk, tryptone, glucose and glycerine (STGG) storage media and analysed for the presence of six bacterial species: *S. pneumoniae, H. influenzae, M. catarrhalis, S. aureus, P. aeruginosa* and *N. meningitidis* using standard microbiological identification techniques (methodology described in Supplementary Methods 1).

**Figure 1.**
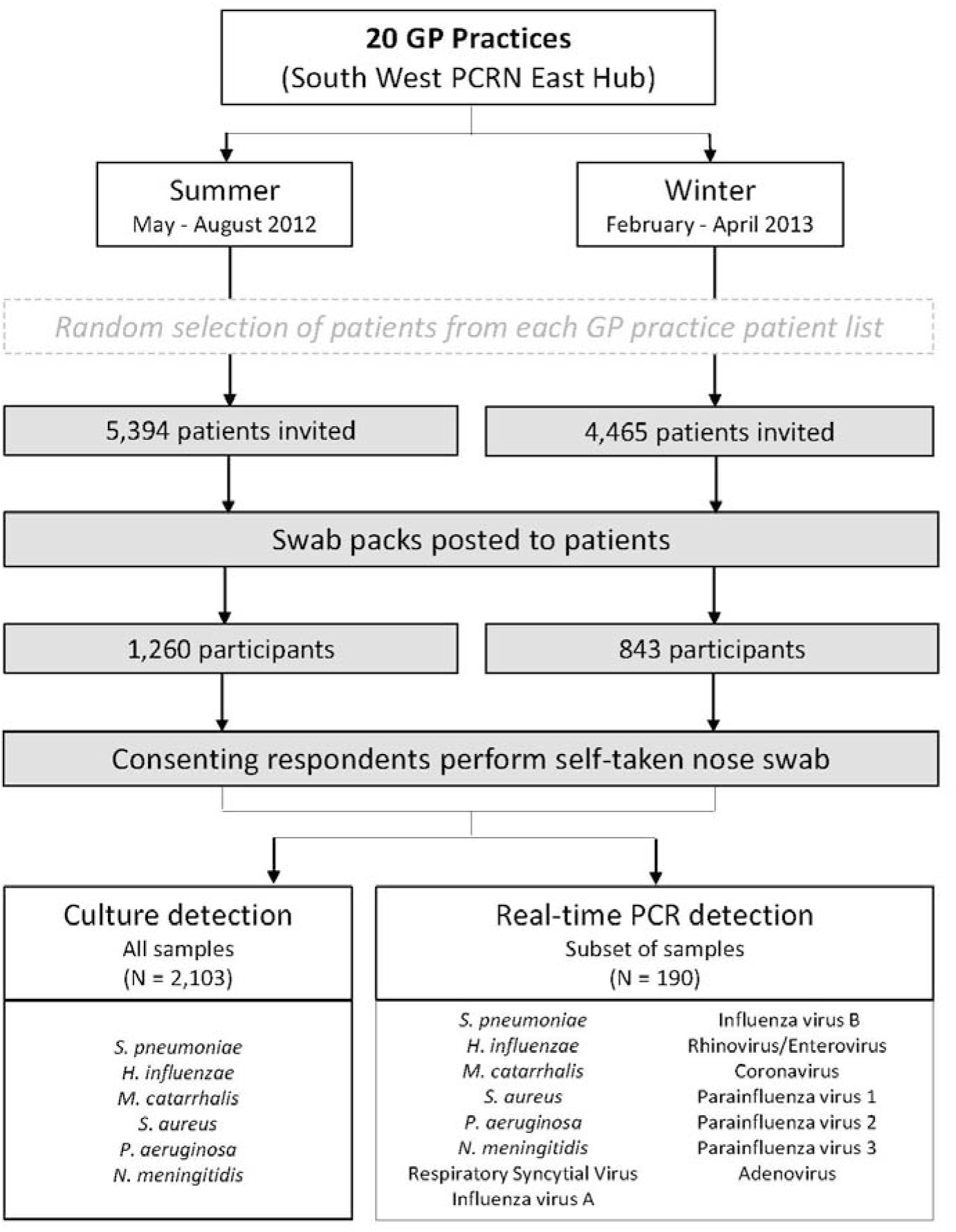
Study design. GP = general practitioner.

### Statistical Analysis

Binary logistic regression (SPSS, version 20) determined the effect of patient characteristics and demographic factors on carriage of the target bacterial species. Univariate linear regression was used to pre-select variables for inclusion in the final model (at Type-I threshold *α* = 0.10). Multivariable binomial logistic regression was then performed using these pre-selected variables, followed by backward elimination of those not meeting a Type-I threshold *α* = 0.05 within the final model. Crude and adjusted odds ratios (OR), 95% confidence intervals and p-values were recorded. Final model fit was assessed using Nagelkerke pseudo-R^2^ values and Receiver Operating Characteristic (ROC) analysis for calculation of the area under the curve (AUC). High pseudo-R^2^ values and an AUC significantly different from 0.5 indicate a model that predicts carriage better than chance variation.

Variables assessed for inclusion by univariate analyses included age (in years), recent RTI (self-reported, RTI within a month prior to swabbing), geographical location (practices were grouped into six geographical areas, recent use of antibiotics (self-reported, antibiotics taken within a month prior to swabbing), immunisation status (self-reported, immunisations in-line with the national schedule), season (winter or summer swabbing) and presence of other microbial species within the same sample.

### Phenotypic and Epidemiological Typing

Phenotypic and epidemiology typing of *S. pneumoniae, H. influenzae* and *S. aureus* was performed in order to determine serotype, sequence type, antibiotic resistance gene and vaccine candidate gene distribution among isolates. PCR serotyping was used to determine serotypes of *S. pneumoniae* and *H. influenzae* identified within summer samples only. *In silico* serotyping was used to determine serotypes of *S. pneumoniae* and *H. influenzae* identified within summer and winter samples. Methicillin resistance typing was done on *S. aureus* isolates identified within both summer and winter samples to determine the prevalence of methicillin-resistant *S. aureus* (MRSA).

#### Nucleic acid extraction

For serotyping and whole genome sequencing, genomic DNA was extracted from bacterial colonies using the QIAamp DNA Mini Kit Blood (Qiagen, Germany) according to the manufacturer’s protocol. For real-time PCR, nucleic acid extraction (DNA and RNA) from STGG samples was undertaken using the QIAxtractor automated nucleic acid purification instrument and QIAamp Viral RNA Mini kits (Qiagen, Germany) according to the manufacturer’s vaccum protocol.

#### PCR Serotyping

*S. pneumoniae* isolates were serotyped using the Centre for Disease Control and Prevention (CDC) protocol (9). This involved eight separate PCR reaction pools covering 40 of the most common pneumococcal serotypes (Supplementary Table 1). Each PCR was undertaken in a 12.5μl volume containing 0.5μl 50mM MgCl_2_, 6.25μl 2× Red PCR, 0.0125μl 100mM *cpsA* (capsular polysaccharide biosynthesis gene for identification of *S. pneumoniae*) forward and reverse primers, 100mM serotype forward and reverse primers (volumes in Supplementary Table 1), Ultra-High Quality (UHQ) water and 1μl extracted DNA (or control). Thermal cycling was performed under the following conditions: 94°C for 4 minutes followed by 30 cycles of 94°C for 45 seconds, 54°C for 45 seconds and 72°C for 2 minutes 30 seconds. Serotyping of *H. influenzae* was undertaken using previously published methods (23, 24). Speciation PCR was undertaken for detection of the *ompP2* gene. Each PCR was undertaken in a 12.5μl volume containing 0.625μl 10mM forward and reverse primers, 6.25μl 2× Red PCR Mix, 4μl UHQ water and 1μl extracted DNA (or control). Thermal cycling was performed under the following conditions: 95°C for 2 minutes then 25 cycles of 94°C for 1 minute, 52°C for 1 minute and 72°C for 1 minute and then a final step of 72°C for 8 minutes. Encapsulation PCR was undertaken for detection of the *bexB* gene. Each PCR was undertaken in a 20μl volume containing 1μl 10mM forward and reverse primers, 10μl 2× Red PCR Mix, 6.4μl UHQ water and 1.6μl extracted DNA (or control). Thermal cycling was performed under the following conditions: 95°C for 4 minutes then 30 cycles of 95°C for 30 seconds, 54°C for 30 seconds and 72°C for 45 seconds and then a final step of 72°C for 8 minutes. Serotyping PCR was undertaken for detection of serotype a-f genes. Each PCR was undertaken in a 12.5μl volume containing 0.625μl 10mM forward and reverse primers for each serotype, 6.25μl 2× Red PCR Mix, 4μl UHQ water and 1μl extracted DNA (or control). Thermal cycling was performed under the following conditions: 95°C for 2 minutes then 25 cycles of 94°C for 1 minute, 52°C for 1 minute and 72°C for 1 minute and then a final step of 72°C for 8 minutes. All PCR products were visualised using a 2% high-resolution agarose gel.

#### Whole Genome Sequencing

Sequencing libraries were prepared using Nextera XT (Illumina, USA). Whole genome sequencing was performed using the Illumina MiSeq paired-end chemistry (2×150bp or 2×250bp). Adapter sequences and low quality regions were removed using Trimmomatic (Usadel Lab, 0.32), quality control of reads was performed using FastQC (Babraham Bioinformatics, version 0.11.2) and then assembled *de novo* using Velvet (Victorian Bioinformatics Consortium, version 2.2.5) (25). Assemblies were then improved using AssemblyImprovement (Wellcome Trust Sanger Institute, version 1.140300).

#### In silico serotyping

*In silico* PCR analysis was then undertaken using iPCRess (European Bioinformatics Institute, Exonerate package, version 1.2.0). Primer sequences for *in silico* serotyping used in iPCRess are shown in Supplementary Table 2.

#### Gene Detection

SRST2 (Short Read Sequence Typing, version 0.1.4) (26) was used to screen the whole genome sequences of the selected *S. pneumoniae* and *H. influenzae* isolates for defined sets of genes: housekeeping genes for determination of sequence type (ST), antibiotic resistance genes (ResFinder/Comprehensive Antibiotic Resistance Database (27, 28)) and genes associated with virulence (Pathogenic Bacteria Database (VFBD) (29, 30)). Antigens that are or have been considered as vaccine candidates were of particular interest. *S. pneumoniae* vaccine candidates included *piaA, pspA, pspC/cbpA, psaA, nanA, lytA* and *ply* (31–34). *H. influenzae* vaccine candidates included *hpd, ompP2, ompP5, tbpA/B, hmw1/2* and *Hia* (35, 36). iPCRess was also used to identify penicillin binding protein genes pbp1a, pbp2x and pbp2b genes within *S. pneumoniae* isolates (37) and the *H. influenzae* protein D (*hdp*) gene (38).

Simpson’s Index of Diversity, 1 – D, was used to quantify the diversity of species’ serotypes and STs. The value of *D* gives the probability that two randomly-sampled individuals belong to different species. The index was calculated from the sum across all species of the number of individuals of each species:

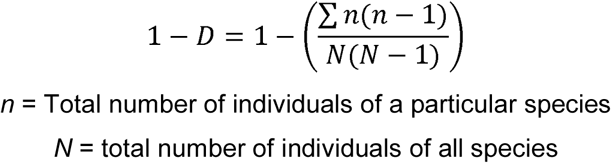

#### MRSA typing

S. *aureus* isolates were tested for methicillin resistance by plating out 10μl of pure culture stored in STGG onto Brilliance MRSA Agar (Oxoid, PO1162) and incubating at 37°C for 18-20 hours. The presence of denim blue colonies of the plates was indicative of MRSA.

### Real-time PCR

A subset of nose swabs (N=190) was analysed by real-time PCR to detect the presence of *S. pneumoniae, H. influenzae, M. catarrhalis, S. aureus, P. aeruginosa, Neisseria meningitidis*, Respiratory Syncytial Virus, Influenza viruses A and B, Rhinovirus/Enterovirus, Coronavirus, Parainfluenza viruses 1-3 and Adenovirus. Details of primers and probes used in the real-time PCR reaction are listed in Supplementary Table 3. Argene PCR kits were used for the detection of Coronavirus and Parainfluenza Virus (71-045) as well as Enterovirus/Rhinovirus (71-042). Bacterial real-time PCR reactions were performed in monoplex 20μl reactions, consisting of 5μl of the primer/probe working stock solution, 10μl Roche Lightcycler 480 Probes Master and 5μl extracted DNA. Viral real-time PCR reactions were done in multiplex 25μl reactions. The Influenza A-B and BMV reaction consisted of 12.5μl Superscript III One-Step RT-PCR System 2× Reaction Mix, 0.8μl Invitrogen SuperScript III RT/Platinum Taq Enzyme Mix, 6.7μl of the primer/probe working stock solution and 5μl extracted RNA. The RSVA-B and MPV reaction consisted of 12.5μl Superscript III One-Step RT-PCR System 2× Reaction Mix, 0.5μl SuperScript III RT/Platinum Taq Enzyme Mix, 6.7μl of the primer/probe working stock solution and 5μl extracted RNA. The Adenovirus reaction consisted of 12.5μl Applied Biosystems TaqMan 2x Universal PCR Mastermix, 7.5 μl of the primer/probe working stock solution and 5μl extracted DNA. For the commercial kits, a 1 in 10 dilution of the reverse transcriptase (RT) was made up in UHQ water. For each sample, 0.15μl of diluted RT was added to 15μl pre-amplification mix. 15μl of the amplification mix was then aliquoted into the reaction tube along with 10μl of extracted RNA. Each monoplex bacterial real-time PCR was run on the Rotor-Gene Q (Qiagen, Germany) with the following thermal cycling conditions: 95°C for 5 minutes and 50 cycles of 95°C for 15 seconds followed by 60°C for 45 seconds. Each multiplex viral real-time PCR was run on the ABI 7500 (Applied Biosystems, USA) with the following thermal cycling conditions: 50°C for 30 minutes, 95°C for 2 minutes and 50 cycles of 95°C for 15 seconds followed by 60°C for 60 seconds. Both commercial PCR kits were run on the Rotor-Gene Q with the following thermal cycling conditions: 50°C for 5 minutes, 95°C for 15 minutes and 45 cycles of 95°C for 10 seconds, 60°C for 40 seconds and 72°C for 25 seconds. Detection of bacterial and viral species was determined using Cycle Threshold (C_T_) values reported by the Rotor-Gene Q and ABI 7500 system.

#### Quantification of bacterial species

Standard curve methodology was used for quantification of bacterial species. Reference strains of *S. pneumoniae* (ATCC 49619), *S. aureus* (clinical isolate), *H. influenzae* (NCTC 11931), *M. catarrhalis* (clinical isolate), *P. aeruginosa* (NCTC 10662) and *N. meningitidis* (MC58) were used to make 1-2 McFarland solutions in 1ml UHQ water. Ten-fold series dilution of each bacterial solution were prepared up to 10^−9^ and plated out in triplicate 10μl spots onto CBA (or chocolated CBA for *H. influenzae*). Following incubation at 37°C for 24 hours (with 5% CO_2_ for S. *pneumoniae, H. influenzae, M. catarrhalis* and *N. meningitidis* plates were counted and CFU/ml values calculated for each dilution. All dilutions were also extracted using the QIAxtractor nucleic extraction and amplified in duplicate on the Rotor-Gene Q. The CFU/ml for each dilution was then correlated to a given C_T_ value to create a standard curve for each species, which was then used for quantification.

### Ecological Analysis

All ecological analyses used real-time PCR results for bacterial species from nose swabs (*N* = 190 species).

#### Nestedness

Nestedness is a measure of organisation within an ecological system that quantifies the ordering of species incidence across space or time. A strongly nested system has a highly predictable sequence of species incidence amongst samples, while a weakly nested system has a more random turnover of species incidence. A nestedness tool, PlotTemp.R, was used within the R graphical user interface (CRAN, version 3.0.1) to determine spatial ordering amongst nose-swab samples analysed with real-time PCR. Real-time PCR datasets were converted to incidence matrices of the presence/absence of each species in each sample. The plot_temp script shuffles each matrix to rank its rows of samples by their richness (most species rich at top) and its columns of species by their incidence (most frequently present to left). Within the final shuffled matrix, presences are represented by red squares and absences by white squares. Their locations in the matrix are compared to a line of perfect fill that would encompass all presences if they were perfectly packed towards the top-left of the matrix. This line then reveals the locations of ‘surprise’ presences lying to its right and ‘surprise’ absences lying to its left. The degree of disorder in the matrix is reported as a matrix ‘temperature’, ranging from 0-100°, calculated as a function of the sum across all surprises of squared deviations from the line of perfect fill. Low temperatures reveal strong nesting, characteristic of highly ordered systems, and high temperatures reveal weak nesting, characteristic of disordered systems. Nestedness was compared between groups of individuals partitioned by age, recent RTI and season.

#### Species distributions

The distributions of the different bacterial and viral species across swab samples were analysed by plotting the observed frequencies of 1, 2, 3, … species within swab samples. This observed distribution was compared to the Poisson distribution as a null model of random species affiliations. Deviations of observed frequencies from Poisson expectation were detected with *p* < 0.05 in a *X*^2^ goodness-of-fit test. Narrower and more peaked distributions than Poisson expectation indicated a more regular than random distribution, with many swab samples containing a similar number of species, providing indirect evidence of mutual repulsion in competition. Wider and flatter distributions than Poisson expectation indicated a more clumped than random distribution, with many swab samples containing few or numerous species, providing indirect evidence of mutual attraction in facilitation. Frequency distributions were compared between groups of individuals partitioned by age, recent RTI and season.

#### Community assembly patterns

Plots of ranked species abundances, measured as bacterial concentrations in CFU/ml, were evaluated against McArthur’s Broken Stick model of neutral community assembly. This model compares an ecological environment to a stick that is broken at random points to produce fragments of lengths that represent the abundances of different species (39, 40). Deviations from neutral expectation were detected with *p* < 0.05 in a *X*^2^ goodness-of-fit test. Observations of rare species below neutral expectation indicate niche characteristics of a dominance hierarchy, while observations of rare species above neutral expectation indicate niche characteristics of resource segregation (15).

## RESULTS

### Participants

A total of 2,103 patients from 20 GP practices across the South East hub of the Wessex Primary Care Research Network participated in self-swabbing across the two time-points. Different individuals were swabbed within each time-point. The characteristics of the study participants are shown in Table 1. Study participants were characterised as having a mean age of 38.7 years (SD 29.4), 8.6% with recent antibiotic use, 34.8% with recent RTI and 80.9% with up-to-date vaccination status. Participation was 23.4% (*n* = 1,260, 95% CI 22.27% to 24.53%) in summer and 18.9% (*n* = 843, 95% CI 17.7% to 20.0%) in winter. Individual GP practice participation varied from 9.3% (*n* = 27) to 33.1% (*n* = 96) in the summer self-swabbing group and 9.1% (*n* = 21) to 30.4% (*n* = 80) in the winter self-swabbing group.

**Table 1.** Characteristics of study participants

|  | **Summer**<br>(N = 1,260) | **Winter**<br>(N = 843) | **Total**<br>(N = 2,103) |
| --- | --- | --- | --- |
| **Age (years), n (%)** |  |  |  |
| Mean (SD) | 37.4 (29.1) | 40.7 (29.8) | 38.7 (29.4) |
| 0-4 | 329 (26.1) | 168 (19.9) | 497 (23.6) |
| 5-17 | 137 (10.9) | 127 (15.1) | 264 (12.6) |
| 18-64 | 465 (36.9) | 286 (33.9) | 751 (35.7) |
| 65+ | 311 (24.7) | 244 (28.9) | 555 (26.4) |
| Missing | 18 (1.4) | 18 (2.1) | 36 (1.7) |
| **Recent antibiotics use, n (%)** |  |  |  |
| Yes | 101 (8.0) | 79 (9.4) | 180 (8.6) |
| No | 1124 (89.2) | 743 (88.1) | 1867 (88.8) |
| Missing | 35 (2.8) | 21 (2.5) | 56 (2.7) |
| **Recent RTI, n (%)** |  |  |  |
| Yes | 365 (29.0) | 367 (43.5) | 732 (34.8) |
| No | 860 (68.3) | 453 (53.7) | 1313 (62.4) |
| Missing | 35 (2.8) | 23 (2.7) | 58 (2.8) |
| **Vaccination Status, n (%)** |  |  |  |
| Up-to-date | 1022 (81.1) | 680 (80.7) | 1702 (80.9) |
| Not up-to-date | 40 (3.2) | 33 (3.9) | 73 (3.5) |
| Missing | 198 (15.7) | 130 (15.4) | 328 (15.6) |

SD = standard deviation, RTI = respiratory tract infection, recent is having occurred with one month of swab sample collection (indicated on the questionnaire).

### Carriage of Bacterial and Viral Species

Unadjusted and age group-specific prevalence of bacterial and viral species carriage within nose swabs detected by culture and real-time PCR are shown in Tables 2–4. All 2,103 nose samples were analysed by culture and a subset of 190 nose samples were analysed by real-time PCR. Carriage was higher in younger age-groups (0-4 and 5-17) for *S. pneumoniae, M. catarrhalis* and *H. influenzae.* Conversely higher carriage levels were observed for *S. aureus* in the older age groups. Comparatively fewer incidences of *P. aeruginosa* or *N. meningitidis* carriage were observed (Tables 2 and 3). Viral carriage was shown to be dominated by Rhinovirus/Enterovirus (13.2%), Coronavirus (10%) and RSV (6.8%) (Table 4). Carriage of bacterial and viral species within the respiratory tract was shown to vary with participant age (*S. pneumoniae* aOR=0.943; *H. influenzae* aOR=0.948; *M. catarrhalis* aOR=0.981, RV/EV FET p< 0.001), recent RTI (*S. pneumoniae* aOR=2.071; RV/EV FET *p*<0.001; COV FET *p*=0.013) and the presence of other species (Table 5). Modelling of viral species prevalence was not possible due to small numbers of viruses detected within this sample set, hence only descriptive statistics are shown in Table 6 for RSV, adenovirus, rhinovirus/enterovirus and coronavirus.

**Table 2.** Carriage of bacterial species in nose swabs detected by culture, stratified by age group

| Carriage, *n* (%) | Age Group (years) | Summer<br>(*N* = 1,260) | Winter<br>(*N* = 843) | Total<br>(*N* = 2,103) |
| --- | --- | --- | --- | --- |
| *S. pneumoniae* | All | 138 (11.0) | 74 (8.8) | 212 (10.1) |
|  | 0-4 | 108 (32.8) | 51 (30.4) | 159 (32.0) |
|  | 5-17 | 18 (13.1) | 18 (14.4) | 36 (13.7) |
|  | 18-64 | 5 (1.1) | 2 (0.7) | 7 (0.9) |
|  | ≥65 | 6 (2.0) | 2 (0.8) | 8 (1.5) |
| *M. catarrhalis* | All | 31 (2.5) | 64 (7.6) | 95 (4.5) |
|  | 0-4 | 19 (5.8) | 31 (18.5) | 50 (10.1) |
|  | 5-17 | 1 (0.7) | 15 (12.0) | 16 (6.1) |
|  | 18-64 | 7 (1.5) | 6 (2.1) | 13 (1.7) |
|  | ≥65 | 4 (1.3) | 9 (3.7) | 13 (2.4) |
| *H. influenzae* | All | 34 (2.7) | 45 (5.4) | 79 (3.8) |
|  | 0-4 | 24 (7.3) | 31 (18.5) | 55 (11.1) |
|  | 5-17 | 7 (5.1) | 10 (8.0) | 17 (6.5) |
|  | 18-64 | 1 (0.2) | 2 (0.7) | 3 (0.4) |
|  | ≥65 | 2 (0.7) | 0 (0.0) | 2 (0.4) |
| *S. aureus* | All | 269 (21.5) | 149 (17.7) | 418 (19.9) |
|  | 0-4 | 32 (9.7) | 16 (9.5) | 48 (9.7) |
|  | 5-17 | 48 (35.0) | 21 (16.8) | 69 (26.3) |
|  | 18-64 | 115 (24.8) | 68 (23.8) | 183 (24.4) |
|  | ≥65 | 71 (23.2) | 41 (16.8) | 112 (20.4) |
| *P. aeruginosa* | All | 20 (1.6) | 7 (0.8) | 27 (1.3) |
|  | 0-4 | 9 (2.7) | 2 (1.2) | 11 (2.2) |
|  | 5-17 | 1 (0.7) | 1 (0.8) | 2 (0.8) |
|  | 18-64 | 6 (1.3) | 1 (0.3) | 7 (0.9) |
|  | ≥65 | 3 (1.0) | 2 (0.8) | 5 (0.9) |
| *N. meningitidis* | All | 0 (0.0) | 1 (0.1) | 0 (0.0) |
|  | 0-4 | 0 (0.0) | 0 (0.0) | 0 (0.0) |
|  | 5-17 | 0 (0.0) | 0 (0.0) | 0 (0.0) |
|  | 18-64 | 0 (0.0) | 1 (0.3) | 1 (0.1) |
|  | ≥65 | 0 (0.0) | 0 (0.0) | 0 (0.0) |

**Table 3.** Carriage of bacterial in nose swabs detected by real-time PCR, stratified by age group

| **Carriage, n (%)** | **Age Group (years)** | **Summer (N = 115)** | **Winter (N = 75)** | **Total (N = 190)** |
| --- | --- | --- | --- | --- |
| *S. pneumoniae* | All | 40 (34.8) | 24 (32.0) | 64 (33.7) |
|  | 0-4 | 21 (52.5) | 11 (55.0) | 32 (53.3) |
|  | 5-17 | 11 (35.5) | 8 (42.1) | 19 (38.0) |
|  | 18-64 | 2 (9.1) | 3 (16.7) | 5 (12.5) |
|  | ≥65 | 6 (27.3) | 2 (11.1) | 8 (20.0) |
| *M. catarrhalis* | All | 28 (24.3) | 17 (22.7) | 45 (23.7) |
|  | 0-4 | 18 (45.0) | 8 (40.0) | 26 (43.4) |
|  | 5-17 | 6 (19.4) | 4 (21.1) | 10 (20.0) |
|  | 18-64 | 1 (4.5) | 3 (16.7) | 4 (10.0) |
|  | ≥65 | 3 (13.6) | 2 (11.1) | 5 (12.5) |
| *H. influenzae* | All | 34 (29.6) | 33 (44.0) | 67 (35.3) |
|  | 0-4 | 19 (47.5) | 15 (75.0) | 34 (56.7) |
|  | 5-17 | 9 (29.0) | 9 (47.4) | 18 (36.0) |
|  | 18-64 | 2 (9.1) | 6 (33.3) | 8 (20.0) |
|  | ≥65 | 4 (18.2) | 3 (16.7) | 7 (17.5) |
| *S. aureus* | All | 50 (43.5) | 33 (44.0) | 83 (43.7) |
|  | 0-4 | 10 (25.0) | 4 (20.0) | 14 (23.3) |
|  | 5-17 | 17 (54.8) | 8 (42.1) | 25 (50.0) |
|  | 18-64 | 12 (54.5) | 12 (66.7) | 24 (60.0) |
|  | ≥65 | 11 (50.5) | 9 (50.5) | 20 (50.0) |
| *P. aeruginosa* | All | 54 (47.0) | 15 (20.0) | 69 (36.3) |
|  | 0-4 | 17 (42.5) | 1 (5.0) | 18 (30.0) |
|  | 5-17 | 16 (51.6) | 4 (21.1) | 20 (40.0) |
|  | 18-64 | 12 (54.5) | 7 (38.9) | 19 (47.5) |
|  | ≥65 | 9 (40.9) | 3 (16.7) | 12 (30.0) |
| *N. meningitidis* | All | 0 (0.0) | 0 (0.0) | 0 (0.0) |
|  | 0-4 | 0 (0.0) | 0 (0.0) | 0 (0.0) |
|  | 5-17 | 0 (0.0) | 0 (0.0) | 0 (0.0) |
|  | 18-64 | 0 (0.0) | 0 (0.0) | 0 (0.0) |
|  | ≥65 | 0 (0.0) | 0 (0.0) | 0 (0.0) |

**Table 4.** Presence of viral species in nose swabs detected by real-time PCR

| **Presence, *n* (%)** | **Summer**<br>**(*N* = 115)** | **Winter**<br>**(*N* = 75)** | **Total**<br>**(*N* = 190)** |
| --- | --- | --- | --- |
| Influenza virus A | 0 (0.0) | 2 (2.7) | 2 (1.1) |
| Influenza virus B | 0 (0.0) | 0 (0.0) | 0 (0.0) |
| Respiratory syncytial virus | 7 (6.1) | 6 (8.0) | 13 (6.8) |
| Adenovirus | 5 (4.3) | 1 (1.3) | 6 (3.2) |
| Rhinovirus/enterovirus | 19 (16.5) | 6 (8.0) | 25 (13.2) |
| Coronavirus | 12 (10.4) | 7 (9.3) | 19 (10.0) |
| Parainfluenza virus | 0 (0.0) | 1 (1.3) | 1 (0.5) |
| Metapneumovirus | 0 (0.0) | 0 (0.0) | 0 (0.0) |

**Table 5.**
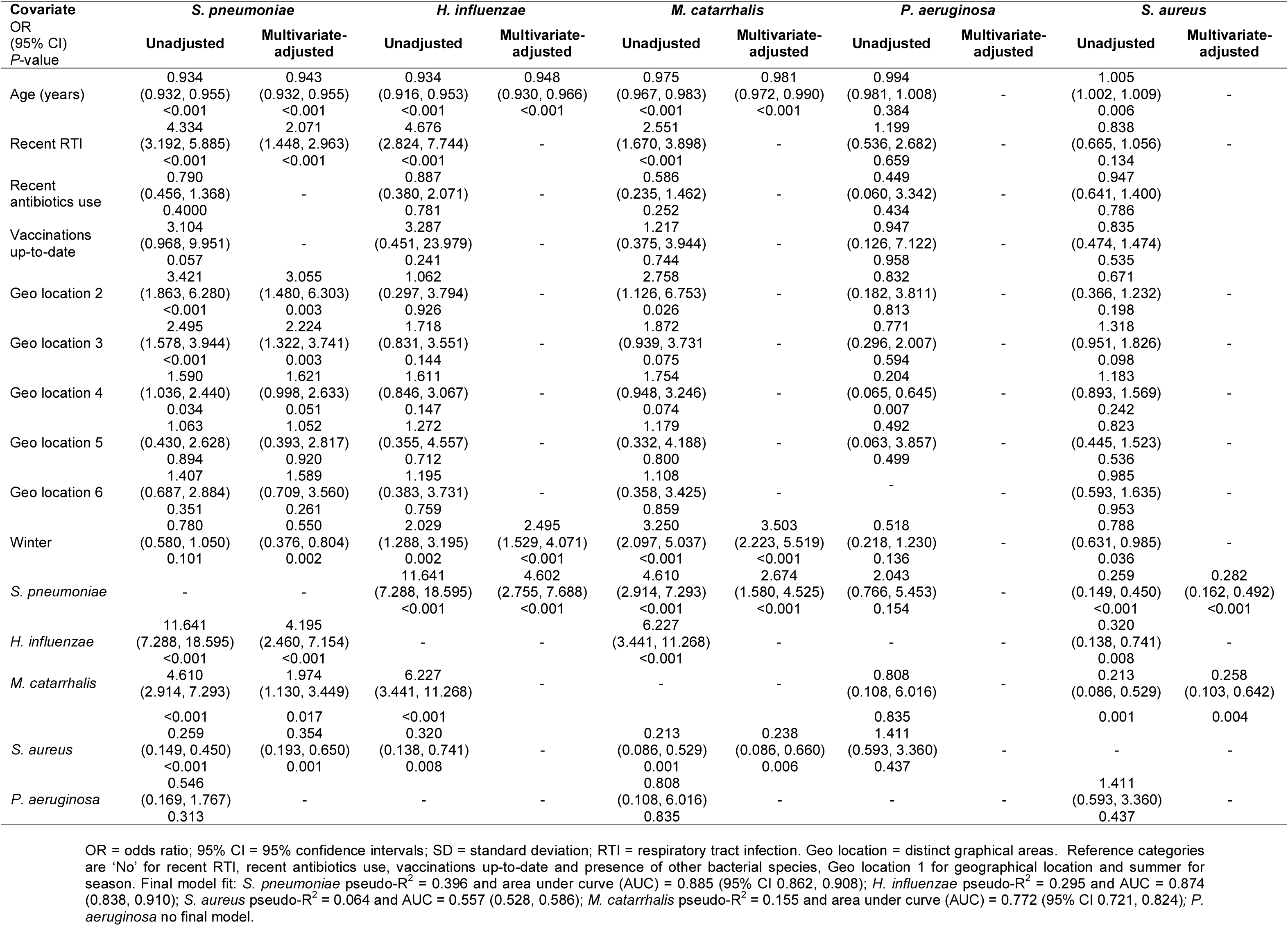
Unadjusted and multivariate-adjusted logistic regression models for nasal carriage of bacterial species

**Table 6.**
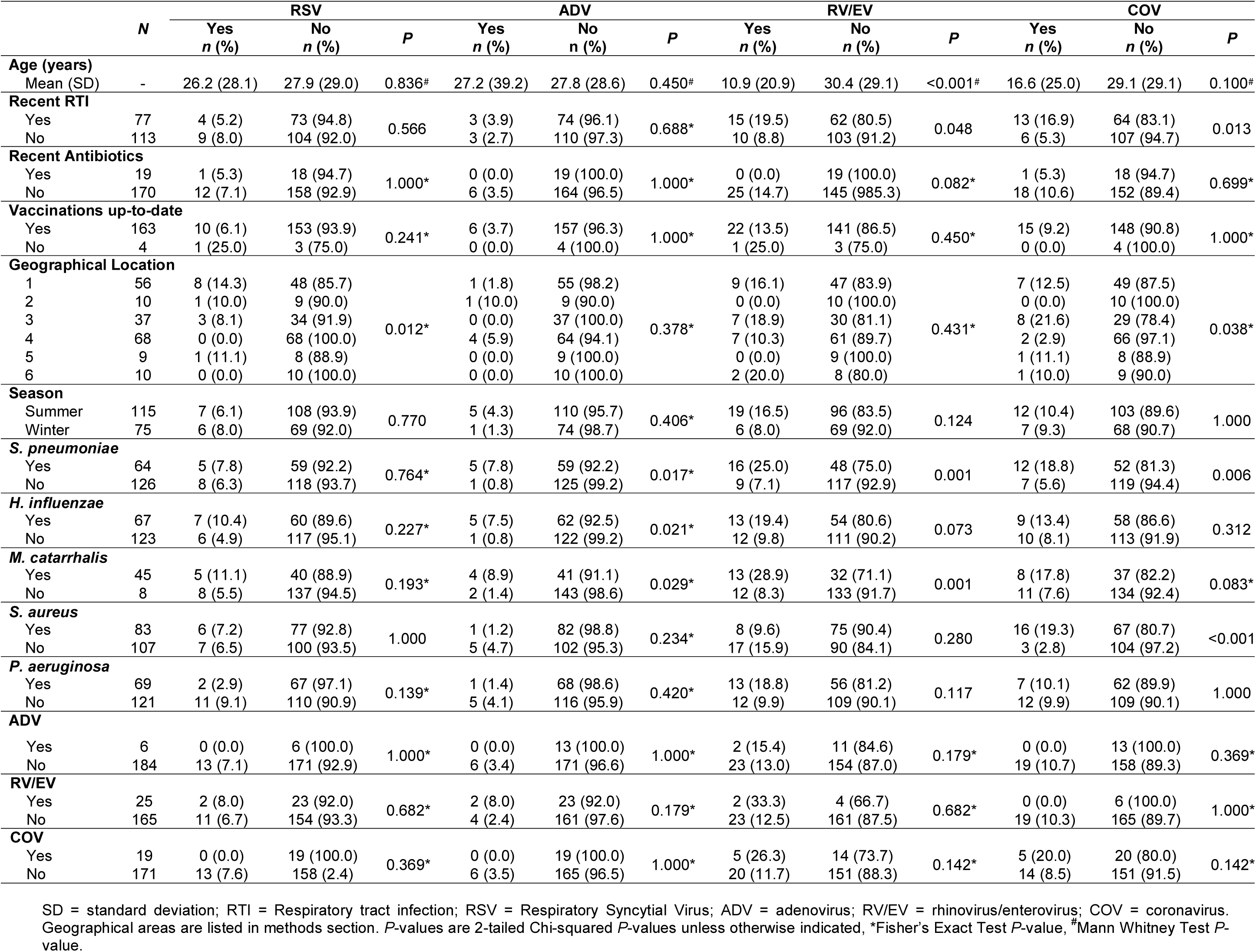
Summary Statistics of Viral Prevalence in Nose Swabs

### Phenotypic and Epidemiological Typing

#### Serotyping

A total of 209 *S. pneumoniae* isolates were serotyped, 10.5% (*n* = 22) of which were PCV7/PCV13 vaccine types, 85.2% (*n* = 178) non-vaccine types and 4.3% (*n* = 9) non-typeable/unknowns. The most common serotypes were 11A/D (11.5%, *n* = 24), 6C (8.6%, *n* = 18), 15B/C (8.6%, *n* = 18), 24A/B/F (6.7%, *n* = 14), 23B (6.7%, *n* = 14), 19A (6.2%, *n* = 13) and 35F/47F (6.2%, *n* = 13). Seventy-four *H. influenzae* isolates were serotyped, all of which were identified as non-typeable *H. influenzae* (NTHi).

#### Sequence Typing

Sixty-three distinct sequence types (ST) of *S. pneumoniae* were identified within the set of isolates. There were also 11 isolates of previously unknown ST. The most common STs were 62 (10.5%, *n* = 22), 199 (10.0%, *n* = 21), 162 (5.7%, *n* = 12), 439 (5.3%, *n* = 11) and 1635 (5.3%, *n* = 11). ST199 was frequently associated with serotypes 19A (9/13, 69.2%) and 15B/C (11/18, 61.1%). ST162 was frequently associated with serotype 24A/B/F (10/14, 71.4%). ST439 was frequently associated with serotype 23B (11/14, 78.6%) ST1635 was frequently associated with serotype 35F/47F (11/13, 84.6%). ST62 was frequently associated with serotype 11A/D (22/24, 91.7%). The set of *S. pneumoniae* STs gave a Simpson’s Index of Diversity (1-D) of 0.96.

Forty-two distinct sequence types of *H. influenzae* were identified within the set of isolates. The *H. influenzae* isolates were diverse with only one or two isolates of each ST identified. The most common STs were 474 (6.8%, *n* = 5), 57 (4.1%, *n* = 3), 348 (4.1%, *n* = 3), 569 (4.1%, *n* = 3) and 1215 (4.1%, *n* = 3). Thirteen isolates were of previously unknown sequence type. The set of *H. influenzae* isolates gave a Simpson’s Index of Diversity (1-D) of 0.99.

#### Antibiotic Resistance

Genes associated with tetracycline resistance, *tetM* (*n* = 9, 4.3%), *tetS* (*n* = 2, 1.0%) and *tet32* (*n* = 1, 0.5%); macrolide resistance *mefA* (*n* = 4, 1.9%) and *msrD* (*n* = 4, 1.9%); erythromycin resistance *ermB* (*n* = 6, 2.9%) and chloramphenicol resistance *cat* (n = 1, 0.5%) were detected in *S. pneumoniae* isolates. *S. pneumoniae* beta-lactam susceptibility genes were *pbp1a* (*n* = 195, 93.3%), *pbp2×* (*n* = 201, 96.2%) and *pbp2b* (*n* =185, 88.5%). Genes associated with beta-lactam resistance *blaTEM* (*n* = 6, 8.1%) and aminoglycoside resistance *aph(3’)-I* (*n* = 3, 4.1%) were identified in *H. influenzae* isolates.

#### Vaccine Candidate Gene Detection

Prevalence of vaccine candidate genes in the 34 *S. pneumoniae* isolates were *lytA* (*n* = 34, 100%), *piaA* (*n* = 34, 100%), *ply* (*n* = 34, 100%), *psaA* (*n* = 34, 100%), *pspA* (*n* = 9, 26.5%), *pspC/cbpA* (*n* = 28, 82.4%) and *nanA* (*n* = 32, 94.1%). Prevalence of vaccine candidate genes in the 32 *H. influenzae* isolates were *hpd* (*n* = 27, 84.4%), *ompP2* (*n* = 32, 100%), *ompP5* (n = 29, 90.6%), *tbpA/B* (*n* = 32, 100%), *hmw1* (*n* = 5, 15.6%), *hmw2* (*n* = 5, 15.6%) and *hia* (*n* = 11, 34.4%).

#### MRSA Typing

A total of 12 (2.9%) methicillin-resistant *S. aureus* isolates were identified within 409 nose isolates of *S. aureus* tested. This included 10 (3.8%) in summer nose swabs and 2 (1.4%) in winter nose swabs.

### Species Organisation and Community Assembly Patterns

#### Nestedness

Microbial species showed a more ordered incidence across samples, indicated by lower nestedness temperature, *T*, amongst individuals with no recent RTI (*T* = 8.47° [Matrix fill = 0.20]) than amongst those with recent RTI (T = 12.54° [Matrix fill = 0.22]). Figure 2 shows the more disordered turnover of species between samples for recently ill individuals (Fig. 2a) compared to healthy individuals (Fig. 2b). Incidence was also more ordered amongst winter samples (*T* = 7.04° [Matrix fill = 0.18]) than summer samples (*T* = 12.39° [Matrix fill = 0.24]), and amongst older individuals (18-64 year olds *T* = 8.92° [Matrix fill = 0.23]; ≥65 year olds *T* = 6.45° [Matrix fill = 0.15]) than younger individuals (0-4 year olds *T* = 11.67° [Matrix fill = 0.27]; 5-17 year olds *T* = 15.21° [Matrix fill = 0.28]).

**Figure 2.**
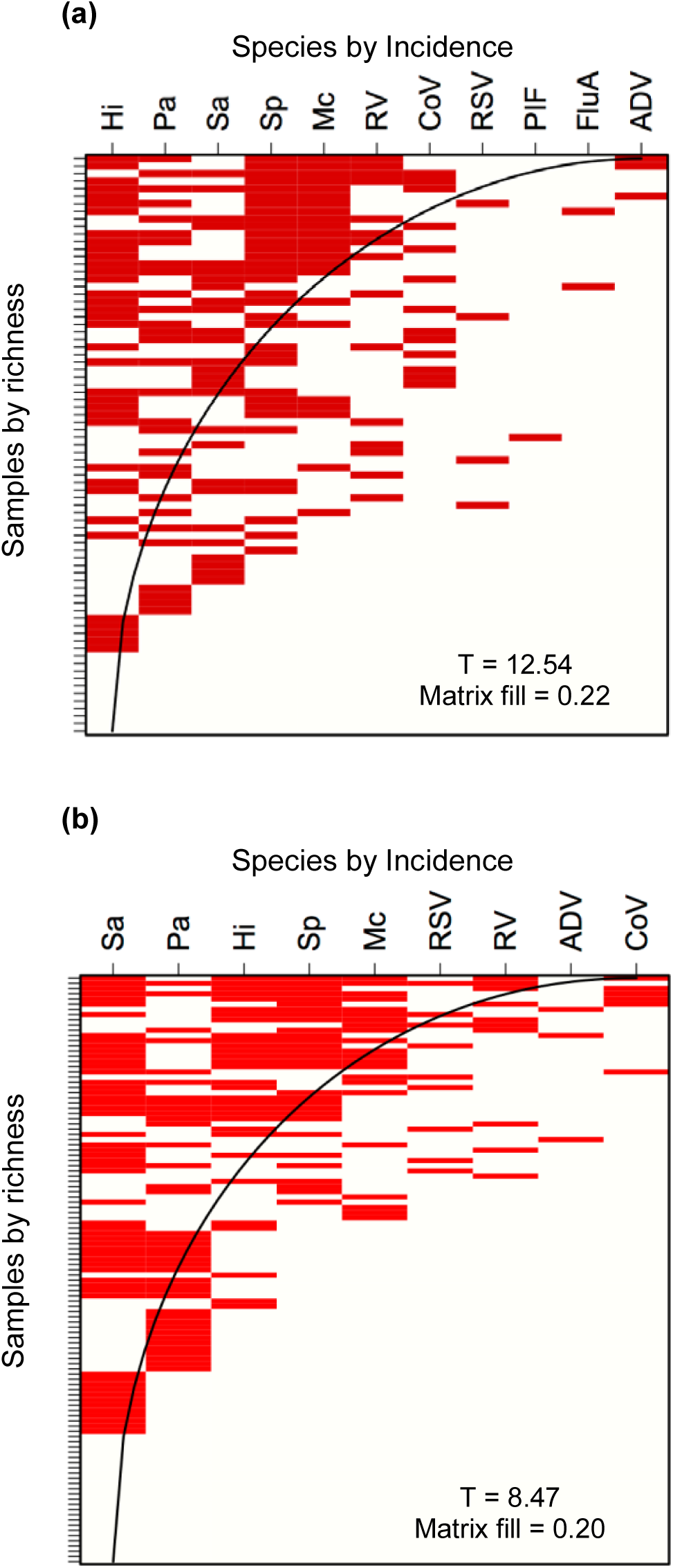
Incidence matrices of nose swab samples detected by real-time PCR in participants (a) with recent RTI (b) with no recent RTI. Red cells show presences, white show absences, curve delineates perfect fill, T = nestedness temperature. Pa = *P.aeruginosa*, Sa = *S. aureus*, Sp = *S. pneumoniae*, Hi = *H. influenzae*, Mc = *M. catarrhalis*, RV = rhinovirus/enterovirus, CoV = coronavirus, RSV = respiratory syncytial virus, ADV = adenovirus, PIF = parainfluenza virus, FluA = influenza virus A.

#### Species Distribution

Frequency distributions of microbial species richness deviated from random amongst summer samples (X^2^ = 21.75, df = 5, *p* < 0.001), 0-4 year olds (X^2^ = 20.42, df = 6, *p* = 0.002) and individuals without recent RTI (X^2^ = 15.44, df = 5, *p* = 0.009). These classifications had wider frequency distributions than Poisson expectation, with more occurrences of few or many species than would be expected from a random distribution. This wider distribution indicates clumping of species. In particular *S. pneumoniae*, *M. catarrhalis* and *H. influenzae* were shown to co-occur (*S. pneumoniae* - *M. catarrhalis* OR 4.160 (2.914-7.293) *p* <0.001; *S. pneumoniae* - *H. influenzae* OR 11.641 (7.288-18.595) *p* <0.001; *H. influenzae* - *M. catarrhalis* OR 6.277 (3.441-11.268) *p* <0.001) (Table 5). Conversely, frequency distributions of microbial species richness did not deviate detectably from random amongst winter swabs samples (X^2^ = 6.61, df = 6, *p* = 0.358), 5-17 year olds (X^2^ = 8.35, df = 5, *p* = 0.138), 18-64 year olds (X^2^ = 5.32, df = 4, *p* = 0.256), ≥65 year olds (X^2^ = 0.153, df = 3, *p* = 0.985) and individuals with recent RTI (X^2^ = 7.46, df = 5, *p* = 0.189). For these classifications we therefore found no evidence of interactions between species, either facilitative or competitive.

#### Community Assembly Patterns

Nose samples deviated from the neutral assembly model for samples from summer (X^2^ = 12.36, df = 2, *p* = 0.015), individuals aged 5-17 years (X^2^ = 11.24, df = 2, *p* = 0.024), 18-64 years (X^2^ = 28.78, df = 2, *p* < 0.001) and ≥65 years (X^2^ = 29.69, df = 2, *p* < 0.001) (Age models shown in Figure 3). Within these classifications, rarer species had lower abundances than neutral expectation, indicative of niche characteristics of a dominance hierarchy (cf. Fig. 3a for the 0-4 year age classification with no difference from neutral, and Fig. 3b-d for the other age classifications all deviating from neutral). No detectable deviations from neutral assembly were apparent for samples from winter (X^2^ = 3.05, df = 2, *p* = 0.549), individuals aged 0-4 years (X^2^ = 1.68, df = 2, *p* = 0.794), individuals with recent RTI (X^2^ = 7.39, df = 2, *p* = 0.117) and individuals without recent RTI (X^2^ = 9.03, df = 2, *p* = 0.05).

**Figure 3.**
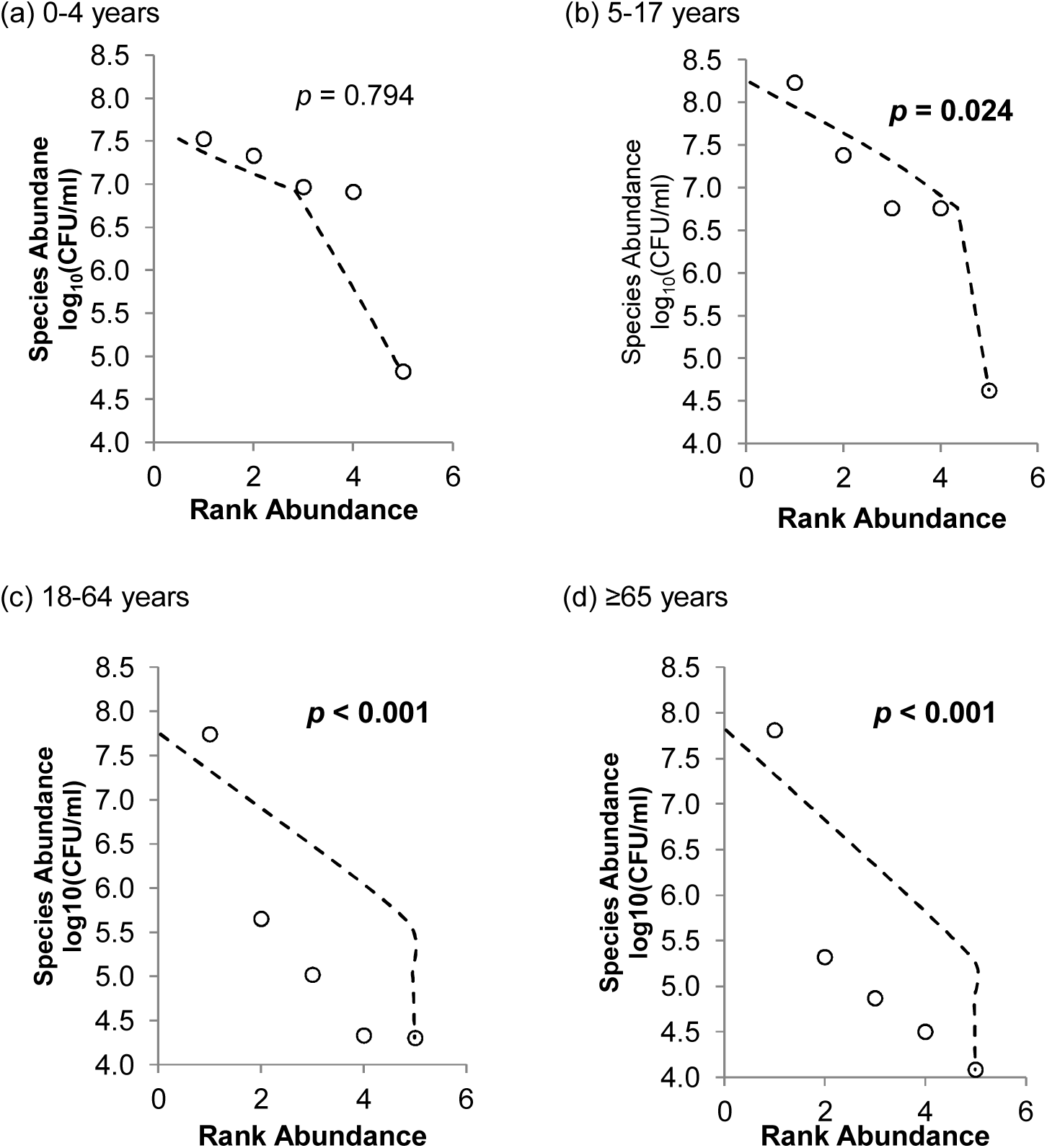
Species abundance curves (CFU/ml) for nose swabs by age group. Dots show species abundance, dashed line is MacArthur’s Broken Stick model of neutral community assembly; deviations with X^2^ *P*-value < 0.05 in bold. CFU = colony-forming unit.

## DISCUSSION

Understanding the prevalence and community distribution of bacterial and viral pathogens, particularly where carriage is a prerequisite for progression to disease, is a key undertaking in the on-going development of interventional responses to changing epidemiology. Here we set out to determine the distribution of pathobiont bacterial and viral species commonly associated with respiratory tract infections using culture-independent approaches and ecological models of community assemblage as well as co-occurrence. Using multivariate analysis of microbial prevalence, we demonstrated that age affects carriage for a number of bacteria and viruses, with young participants experiencing higher carriage of *S. pneumoniae*, *H. influenzae, M. catarrhalis* and rhinovirus/enterovirus. Recent RTI was also associated with increased prevalence of *S. pneumoniae*, rhinovirus/enterovirus and coronavirus. Epidemiological typing revealed few vaccine type *S. pneumoniae* and *H. influenzae* within this set of isolates, which is indicative of the effectiveness of current PCV-13 and Hib vaccines in targeting specific serotypes as well as the effect of herd immunity in protecting individuals within age groups unlikely to have received these vaccines (41, 42). The high proportion of non-vaccine type (NVT) *S. pneumoniae* reflects the process of serotype replacement as a result of vaccination, although it is not thought to lead to future large increases in the incidence of invasive disease caused by NVTs (43). The identification of increasingly common NVT serotypes 6C, 11A/D and 23B and non-typeable *H. influenzae* provides important information for the epidemiological assessment of vaccines and for the development of future vaccines.

The lack of ubiquity of potential vaccine targets within the isolates collected in this study may highlight certain difficulties in the development of protein-based vaccines against these species (32). This knowledge will help to inform the targeting of future treatment and prevention strategies, including the development of new vaccines against *S. aureus* and *M. catarrhalis* (44, 45) and vaccines targeting greater numbers of *S. pneumoniae* serotypes (46). High intra-species diversity within these respiratory isolates is potentially a result of the pressures imposed by current vaccination strategies (47) as well as the high levels of recombination in *S. pneumoniae* (48). Few antibiotic resistance genes within this group of isolates reflects the low and stable levels of antibiotic-resistant respiratory species in the UK (49). Continued monitoring of levels of antibiotic resistance within respiratory isolates within the UK is essential for clinical treatment of infection (49).

The ecological analyses showed a relatively high degree of ordering (nesting) in the incidence of species amongst the respiratory tract communities of older participants, those without recent RTI, and amongst winter samples, in contrast to a more disordered turnover amongst young children, those with recent RTI, and summer samples. Lower levels of nestedness in these individuals may be related to the reduced isolation of species, allowing greater transmission, and the role of the immune system in maintaining species diversity. Furthermore, communities within the respiratory tract were found to have a clumped frequency distribution of species (either few or many species present) amongst young individuals, consistent with facilitative relationships occurring between the species within the upper respiratory tract. Facilitative relationships may enhance transmission opportunities and survival of species carried in the respiratory tract. Respiratory species abundances were more consistent with a dominance hierarchy than a neutral assembly model for older participants, those without recent RTI and summer samples, but not for other classifications. There are likely to be both niche and neutral processes involved in microbial community formation in the upper respiratory tract, with speciation and the environment playing key roles (50, 51). The evidence that niche processes tend towards dominance hierarchies rather than resource segregation suggests that the microbial species compete for similar resources; moreover, communities may have fragile coexistence equilibria, with some dominant competitors functioning as keystone species capable of triggering cascades of extinctions (15). Further development of ecological methodologies will allow predictions of microbial variation as a result of infection, season and increasing age. Monitoring communities in states of health and disease is important for generating knowledge that will be useful in clinical practice (52).

Inferences from this study should be understood in the context of a number of methodological constraints. The characteristics of the 21% (*n* = 2103) of responders may differ from those of non-responders. The study focused on a limited number of species, and did not account for species such as Group A *Streptococci* which may play an important role within this sample site as well as other non-culturable species. Furthermore, the swab samples used were not optimised for the detection of viral species. The questionnaire used for the study was designed for simplicity and ease of completion. This meant that information was not collected on a number of important factors, including smoking status, siblings, and attendance at nursery or day-care facilities. Samples assessed by real-time PCR were limited by the number of species that were targeted as well as the non-random nature of their selection. The ecological models ignore acquired or specific host immunity, which can influence the ecology of the respiratory tract (53). The combined analysis of bacterial and viral species, and the combined analysis of all RTI, may also be simplistic. Neutral theory assumes no variation in the total number of species within a community whilst niche theory ignores dispersal (54, 55). The quantification of nestedness provides only a single measure of matrix temperature per classification of samples, ruling out significance testing.

## Conclusion

The study has provided key epidemiological information on spatio-temporal patterns and trends amongst circulating bacterial species and types. Specifically we have demonstrated for the first time the impact of age and season on the distribution of both bacterial and viral pathobionts, and shown clear inter-genus associations of co-carriage between *S. pneumoniae, M. catarrhalis* and *H. influenzae*. These epidemiological findings on circulating respiratory tract pathogens, that includes the distribution of strains and serotypes amongst different age demographics, underpins our ability to assess risk in these populations and therefore ultimately informs the development of new treatment and prevention strategies as well as targeted and effective antibiotics and vaccination policies.

## Acknowledgments

The authors thank the Bupa Foundation for providing the funding to SCC in order to undertake the study and the Rosetrees Trust for their funding contribution for ALC’s PhD studentship. The authors also thank Shabana Hussain, Beverly Simms, Christine Tumman and Karen Cox for technical assistance throughout the study. The authors thank the Southampton NIHR Wellcome Trust Clinical Research Facility for part-funding SNF. The authors also thank the NIHR Comprehensive Local Research Network (NIHR CLRN), Solent NHS Trust, the South West (East Hub) Primary Care Research Network (PCRN), and Danvers International for their support.

## Author Contributions

ALC contributed to study set-up, data collection, data analysis and writing of the manuscript. CPD and ARK contributed to study-design, data analysis and manuscript preparation. RNW contributed to study set-up, data collection. NB and RA contributed to data collection. DM contributed to study design, set-up and data collection. RA and AT contributed to study design and data collection. SNF, JMJ, HMY, PJR, MAM and MVM contributed to study design, data analysis, proofreading of manuscript. DWC contributed to manuscript writing and proofreading. SCC contributed to study design, data collection, data analysis, and writing and proofreading of the manuscript.

